# A Comprehensive Evaluation of Self Attention for Detecting Regulatory Feature Interactions

**DOI:** 10.1101/2024.08.23.609428

**Authors:** Saira Jabeen, Asa Ben-Hur

**Affiliations:** Colorado State University

## Abstract

The successful use of deep learning in computational biology depends on the ability to extract meaningful biological information from the trained models. Recent work has demonstrated that the attention maps generated by self-attention layers can be interpreted to predict cooperativity between binding of transcription factors, a key feature of gene regulatory networks. We extend this earlier work and demonstrate that the addition of an entropy term yields sparser attention maps that are easier to interpret and provide higher precision interpretations. Furthermore, we performed a comprehensive evaluation of the relative performance of different flavors of attention-based transcription factor cooperativity discovery, and compared methods that use raw attention scores to the use of attribution over the attention scores. Our findings demonstrate the benefit of the entropy-enhanced attention models and provide additional insights that would enable practitioners to make effective use of this valuable tool for biological discovery.

## 1 Introduction

Deep learning has made a major impact in computational biology, addressing complex prediction problems, including the prediction of epigenomic features [1, 2], gene expression [3, 4, 5], deciphering protein-protein interactions [6] and predicting protein structures [7]. In computational biology, deriving biological insight from trained deep networks is critical. However, despite their strength in accurately modeling complex high-dimensional prediction problems, interpreting deep networks remains challenging. Interpretability has been a part of deep learning in computational biology from the very beginning where the authors of DeepBind have introduced a method to interpret the learned weights of the first convolutional layer as a motif model [8]. Another major effort has been the development of attribution methods which aim at discovering the parts of a sequence that most affect the classification results. These include *in-silico* mutagenesis [2] and gradient-based methods [9, 10].

The process of gene expression is regulated by transcription factors (TFs). The regulation of gene expression is orchestrated by the cooperative binding of TFs and other regulatory proteins in close proximity [11, 12]. Identifying these cooperative binding events can reveal insights that are important for understanding specific biological processes [13, 14]. Several methods have been proposed in order to capture regulatory interactions between TFs. The authors of DFIM [15] have proposed a method based on the DeepLIFT attribution method [16]. The more recent SATORI approach leverages the ability of self-attention layers to capture the relationship between different positions in a sequence [17]. We build on this earlier work in several ways:

- We improve model interpretability with the addition of an entropy term that promotes the sparsity of the attention weights, improving both interpretability and precision of the detected regulatory interactions.
- We extend the SATORI framework by scoring regulatory interactions using either raw attention values or attention attribution.
- We perform a comprehensive evaluation of the approach and its variants on simulated and real datasets in the context of deep learning models of varying levels of complexity, demonstrating the effectiveness of attention for this task.

## 2 Related Work

While there has been extensive research on deep learning architectures for genomic data, efforts related to inferring cooperativity between regulatory elements have been somewhat limited. As mentioned above, Greenside et al. [15] introduced a method to identify interactions at both nucleotide and motif resolution levels. Their approach utilizes DeepLIFT by computing the disparity in the attribution scores between a sequence with and without mutation to quantify the relationship between a pair of features. A drawback of DFIM is its significant computational expense and the necessity of a post-processing step. The capability of attention layers to capture relationships between features regardless of their spatial distance is the basis for SATORI [17]. The authors demonstrated comparable results to DFIM with much smaller computational overhead. However, both methods have not been evaluated on realistic simulated datasets that can provide a clear indication of the accuracy of their predictions.

The idea of using attention weights to interpret a trained model has been explored in several other papers. The authors of EPCOT [18] proposed a self-attention-based encoder-decoder architecture to establish a framework for predicting epigenetic state that is able to generalize to conditions that were not a part of their training set. They demonstrated that the learned epigenomic feature embeddings of co-occurring motifs are nearest neighbors in the embedding space, and validated selected interactions using the STRING database. However, the method has not been well tested for the purpose of discovering regulatory interactions. While SATORI uses raw attention values to score regulatory interactions, the ISANREG method performs attribution using integrated gradients over the attention scores to infer TF cooperativity [19]. However, their model is limited to detecting TF cooperativity with a single TF of interest, and is designed specifically for the analysis of TF ChIP-seq datasets. SATORI on the other hand is more general, supporting the detection of TF cooperativity for more complex datasets such as those that probe gene regulation on a global scale. The TIANA model uses an approach similar to SATORI, with the distinction of employing pre-loaded convolutional layers initialized with known motif position-specific score matrices and using attribution scores relative to the attention layers [20]. Their experimental results demonstrate the benefit of this approach, which addresses the challenge in motif matching and filter size choice. However, their model was tested on a limited set of data, and a clearer understanding of the factors that contribute to its performance is still required. The ECHO model takes a very different approach, and uses attribution over Micro-C contacts to identify cooperative binding [21]. This method identifies cooperative binding events without distance limitations, an advantage over sequence-based methods. However, it is limited by the scarcity of Micro-C data.

In conclusion, while several methods have been proposed to infer regulatory interactions and TF cooperativity, existing approaches have limitations. These include computational complexity (DFIM), reliance on scarce data types (ECHO), and the ability to identify interactions only with specific ChIP-seq targeted proteins (ISANREG). Moreover, there is a notable lack of comprehensive evaluation for these methods. This gap in thorough benchmarking limits our understanding of how well these methods perform in realistic scenarios. In this work, we integrate the developments described above into the SATORI framework and provide a detailed benchmark over realistic simulated data to understand the relative performance of different methods of using attention to detect regulatory cooperativity. Furthermore, we demonstrate that adding an entropy penalty term to the attention scores leads to increased network interpretability, especially in noisy real-world datasets.

## 3 Results

### 3.1 SATORI accurately infers TF interactions embedded in simulated datasets

The recently proposed SATORI method provides a general framework for inferring regulatory interactions from a deep learning model that employs a self-attention layer (see Fig. 1). In what follows we compare the performance of various flavors of SATORI using simulated datasets of varying levels of complexity and evaluate the accuracy of the approach. We introduce an extension that incorporates an entropy loss term for the self-attention layer, and demonstrate its ability to increase interpretation accuracy and robustness.

**Figure 1:**
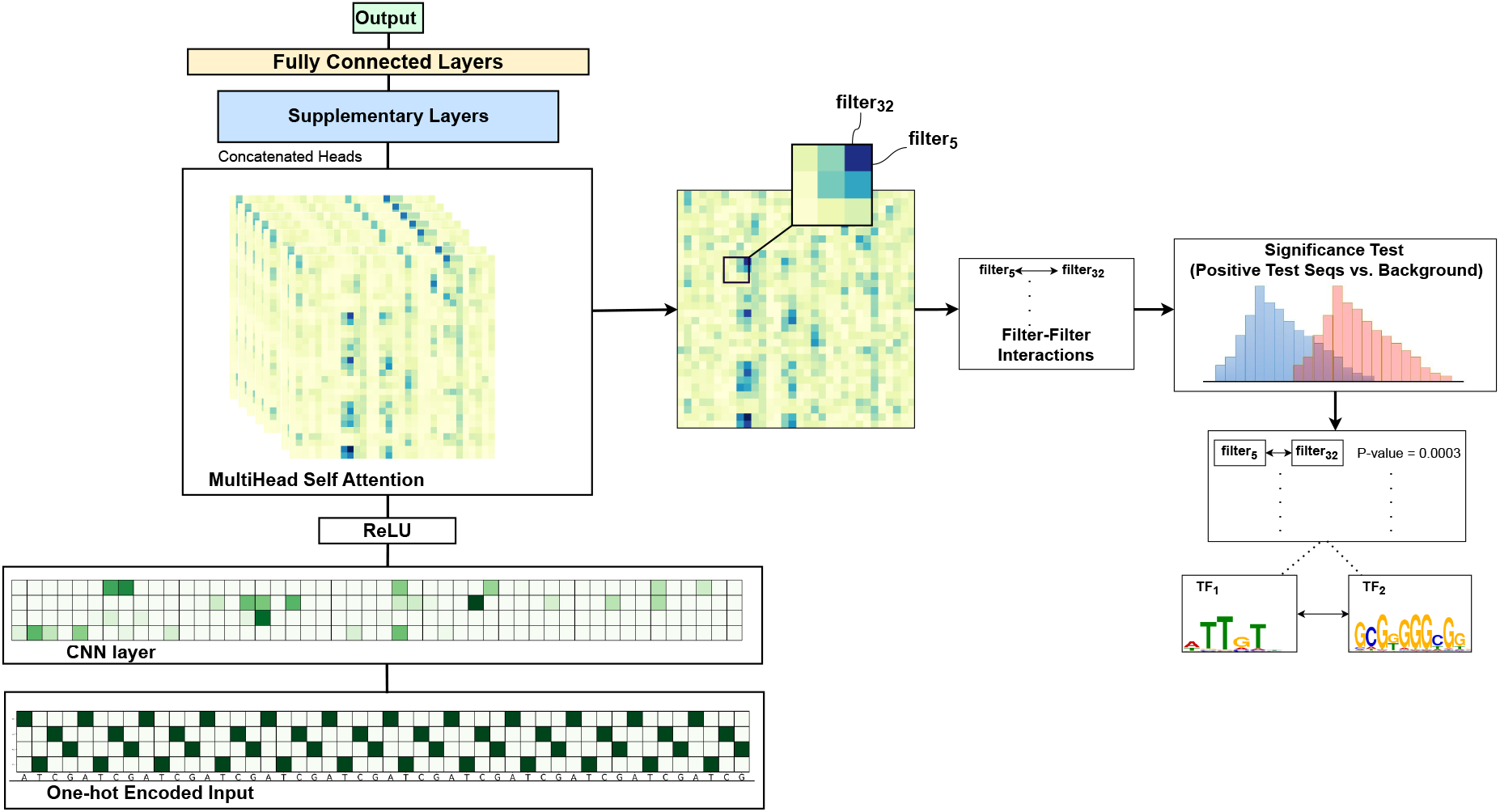
The SATORI framework. The deep learning architecture receives as input a one-hot encoded DNA sequence, which is then processed through a 1D-CNN layer, a self-attention layer, optional supplementary layers and a classification layer. SATORI combines the attention heads and compares the attention values between pairs of filters in positive examples to a background set. It then reports pairs of filters that demonstrate statistical significance.

#### Impact of the Entropy Loss

To comprehensively evaluate the impact of adding an entropy loss term on the quality of predicted interactions, we trained deep learning models with two architectures with varying depth, with and without the entropy loss. Our analysis reveals that the entropy loss maintains the classification accuracy of the models while enhancing precision by promoting sparsity in the attention profile (see Fig. 2). This reduction in false positives aligns with our expectations. We also observe gains in overall accuracy and in most cases it also contributed positively to recall. The marginal decrease in classification accuracy observed in some cases when using the entropy loss indicates a potential loss of information induced by the increased sparsity (see Fig. 2).

**Figure 2:**
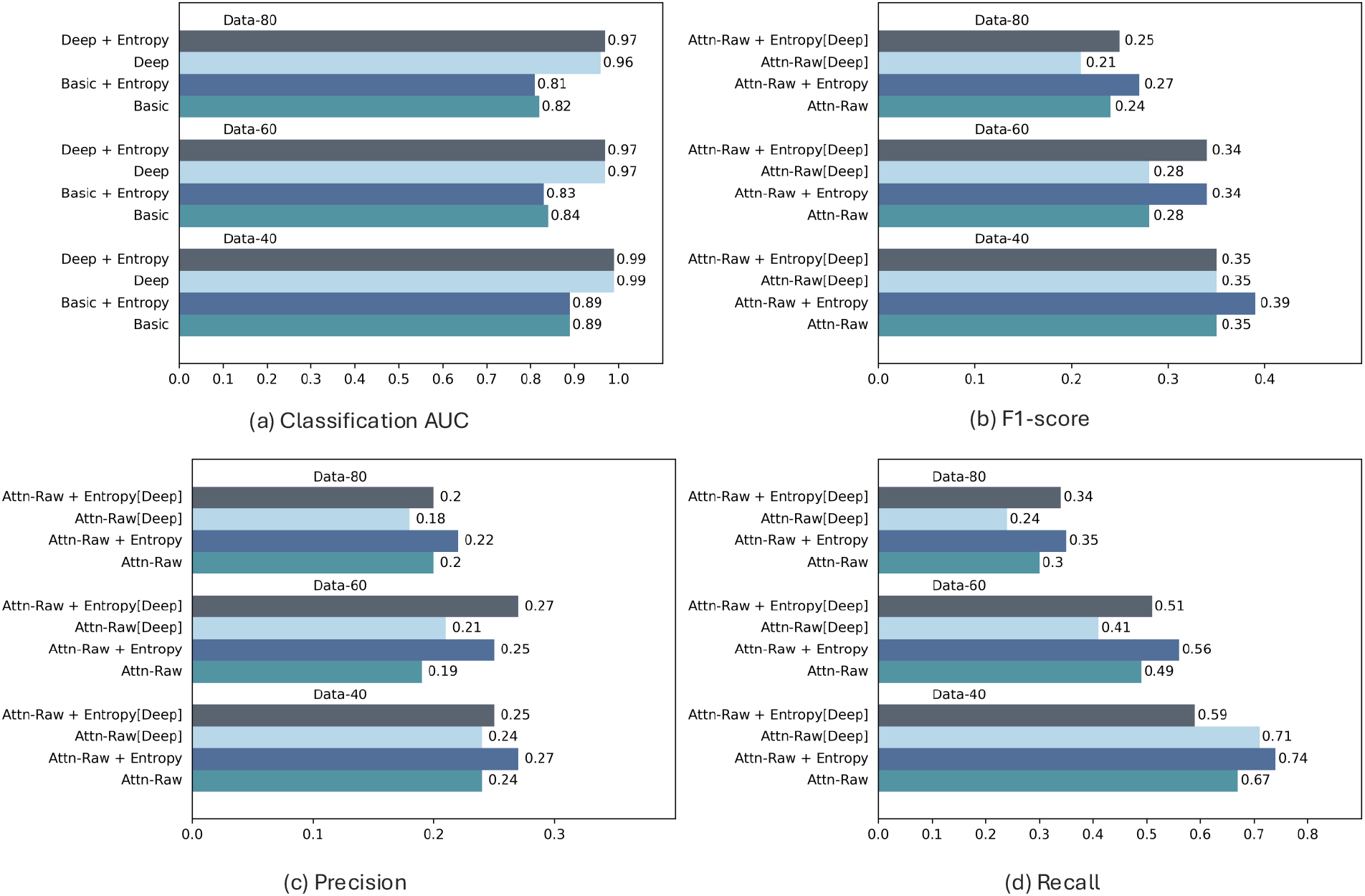
Comparative performance of SATORI on simulated datasets of varying levels of complexity using the Basic and Deep architectures. We show: Classification AUC (a) and interpretation accuracy: F1-score (b), Precision (c) and Recall (d). Results are shown for four models: Basic and Deep trained with and without the entropy loss; results are reported for datasets of increasing levels of complexity, comprising 40, 60, or 80 implanted TF motifs; performance numbers are averages over three instances of each dataset. Results are reported for the version of SATORI that uses raw attention values (Attn-Raw).

We also observe that as the complexity of the simulated dataset increases, both interpretation and classification accuracy decrease, as would be expected. It is also interesting to note that moving from Basic to Deep architecture increases classification AUC, as shown in Fig. 2a. However, interpretation performance demonstrated a slight decrease. This behavior has been consistent across all three simulated datasets.

To obtain a better understanding of the contribution of the entropy term we looked at the distribution of the attention scores with and without entropy (see Fig. 3a). As expected, we observe that the inclusion of entropy results in a distribution of attention scores that is clearly bi-modal. SATORI scores interactions based on the top attention values. The addition of the entropy term has the added benefit of making it trivial to establish a cutoff between attention values that are used for computing feature interactions. The sparsity resulting from the use of entropy can also be observed in the heatmap representing the attention matrix (see Fig. 3b). The vertical lines observed in the heatmap correspond to positions of embedded motifs. Other positions in the sequence attend to these motif positions, suggesting that the model has successfully learned to attend to motif occurrences. In the figure we also observe that the signal in the attention map is more clear in a model trained with an entropy term. Note that the entropy term is multiplied by a regularization parameter whose value needs to be determined. In our experiments, values of the parameter between 0.001 and 0.01 have yielded similar levels of sparsity.

**Figure 3:**
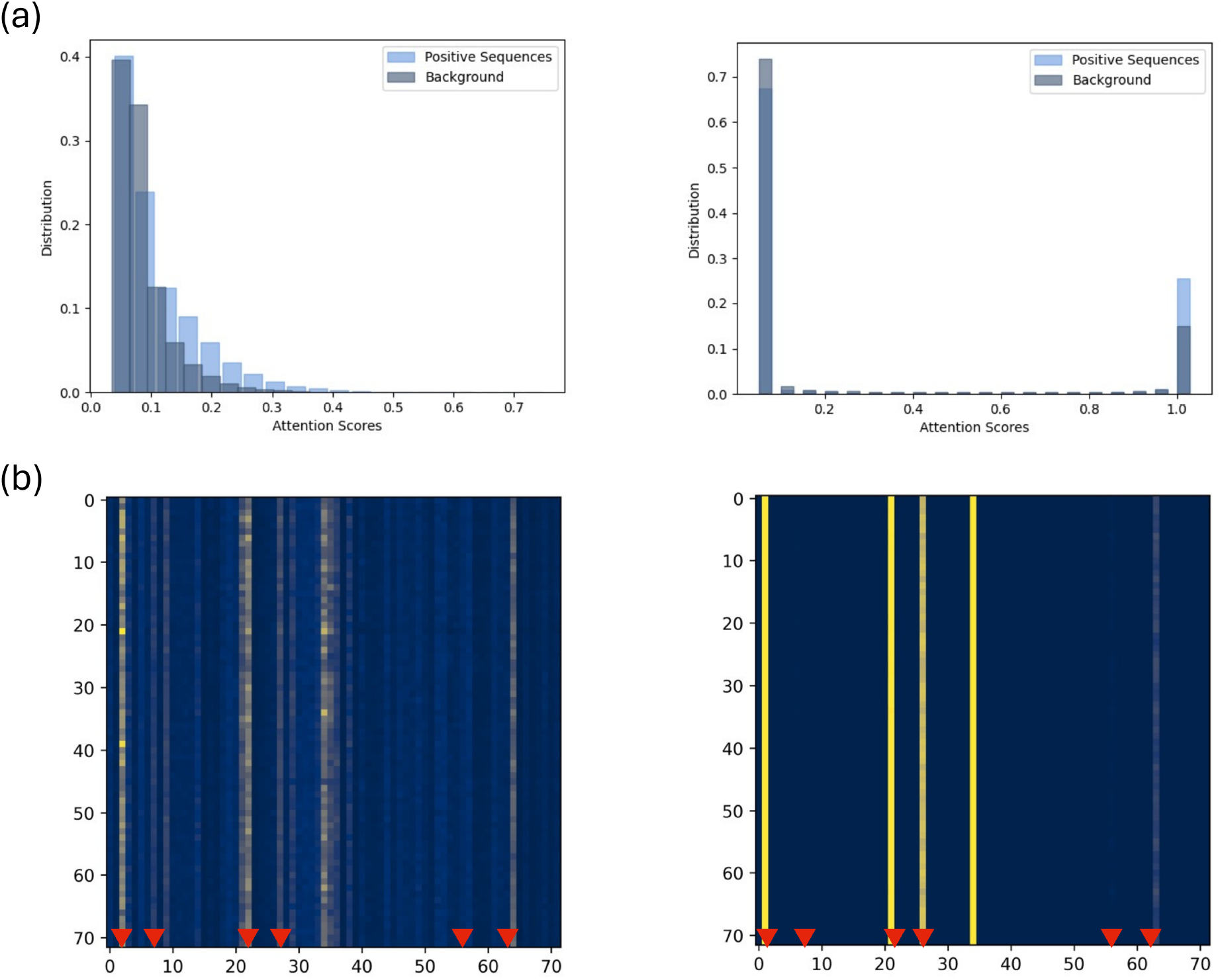
The effect of entropy on attention weights, for a basic model trained without (left) and with (right) the entropy loss using a regularization parameter of 0.01 for Data-80. (a) The distribution of attention weights across accurately predicted test sequences. (b) Heatmaps of attention weights maximized over all attention heads for a true positive sequence. The X-axis and Y-axis in the heatmaps are the positions in the simulated DNA sequence after max pooling. Red arrows indicate the locations of embedded motifs in the original sequence, adjusted for the pooling window.

#### Comparison of raw attention, attention attribution, and DFIM

Here, we compare the interpretation accuracy of SATORI using raw attention scores with that of attention attribution scores and DFIM. In our experiments, we observed that raw attention scores and attention attribution scores give comparable results on simulated datasets as shown in Fig. 4. In fact, the attention attribution matrices exhibit a high similarity to the raw attention matrices (see Fig. S1 in the supplementary material). SATORI with entropy loss outperforms the corresponding model without entropy, both with raw attention scores and attention attribution scores. Both models also outperforms DFIM when used in conjunction with entropy.

**Figure 4:**
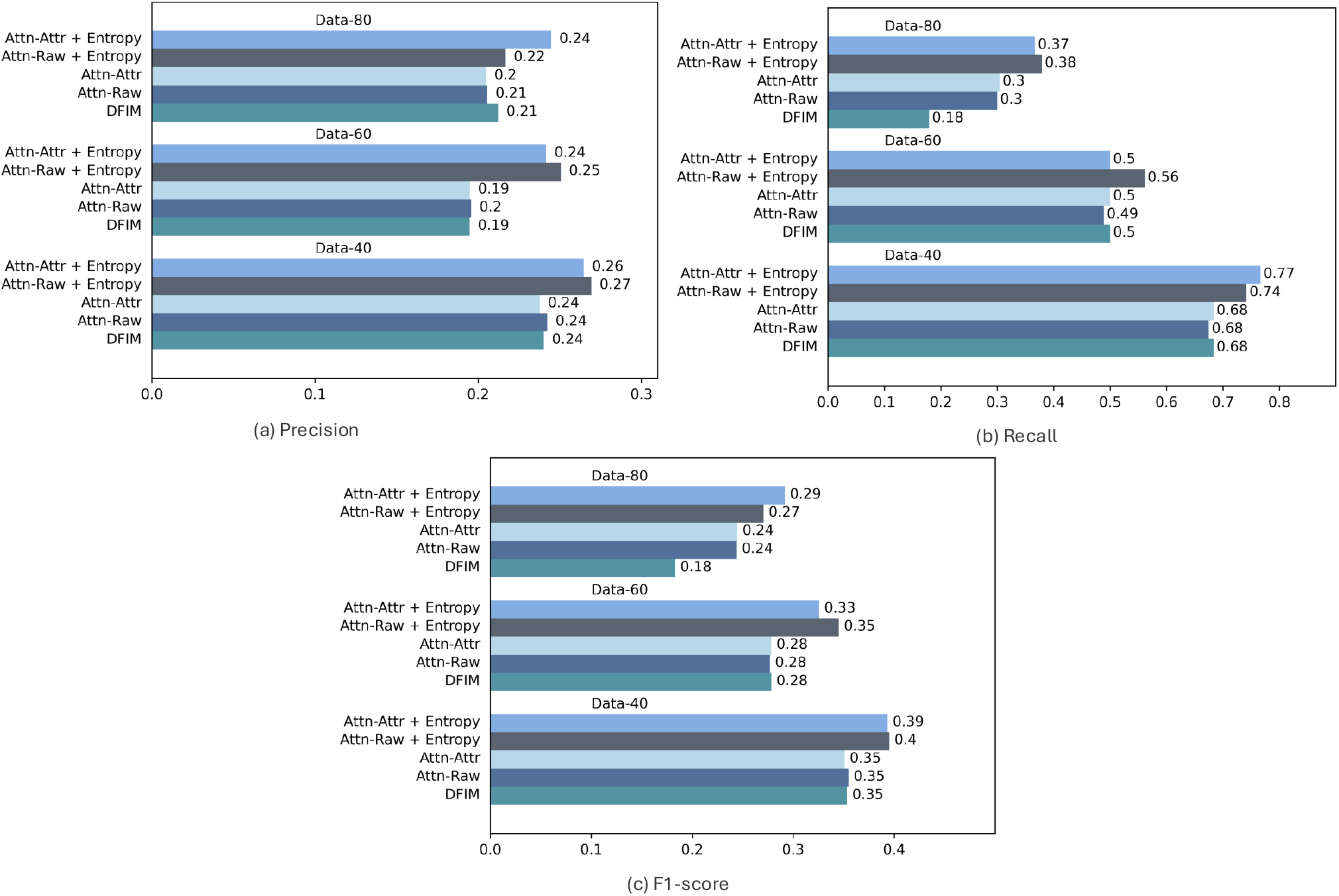
Interpretation accuracy. We compare performance of SATORI using raw attention scores (Attn-Raw), attention attribution (Attn-Attr), and DFIM based on precision (a), recall (b), and F1-score (c). All methods use the same basic architecture; datasets comprise 40, 60, or 80 implanted TF motifs; performance numbers are averages over three instances of each dataset.

An advantage of SATORI with raw attention scores is computation time (see Fig. S2 in the supplement). DFIM and SATORI with attention attribution have the added overhead of requiring the computation of gradients after the model has been trained. For the basic architecture using raw attention scores has only a slight advantage in terms of running time; for the deep architecture gradient computations dominate the running time, and using raw attention scores is around five times faster. The difference with respect to DFIM is even more dramatic, where SATORI with raw attention scores runs over a hundred times faster for the deep architecture.

#### Using pre-loaded filters

The accuracy of inferred interactions depends on the ability of TomTom to accurately match filters with known motifs. The use of pre-loaded CNN filters based on known motifs offers the potential advantage of not requiring computing matches to the trained filters, avoiding a source of potential error. To quantify this, we compared models using trainable CNN filters versus models where the CNN filters were pre-loaded with clustered PSSMs from the Jaspar database, following the setup proposed in [20]. For this purpose, we generated simulated datasets using a sequence length of 1500bp, which will also enable us to demonstrate the ability of self-attention to capture dependencies in longer sequences. We trained both the basic and deep models using hyperparameters selected via random search. Each experiment was repeated on three independently generated dataset instances, and results were averaged to ensure robustness. As expected, using pre-loaded filters leads to improved interpretation accuracy for both the Basic and Deep architectures (Supplementary Fig S4,S5). However, this comes at the expense of a drop in classification accuracy due to reduced flexibility in feature learning (Supplementary Fig S5). We also evaluated the role of positional encoding [20]. While it provides minor improvements in some cases, overall it does not improve classification or interpretation accuracy. We also observe that in some cases attention-attribution failed to match the interpretation accuracy of using raw attention scores, and was somewhat inconsistent in performance. Finally, consistent with our previous results, the deeper model had somewhat lower interpretation accuracy.

#### Incorporating RNNs

In our original work we used a bidirectional recurrent neural network (RNN) layer after the initial CNN layer [17]. Although this design introduces a less direct relationship between the output of the CNN filters and the self-attention layer, we have found that this did not impact interpretability accuracy: on the Data-60 simulated dataset adding an RNN layer (a bidirectional LSTM) to the SATORI-Basic architecture produced an interpretation F1-score of 0.36 compared to 0.33 without the RNN (these represent an average over ten instances). Adding the RNN also improved classification accuracy measured by area under the ROC curve from 0.88 to 0.96. The use of an RNN led to a reduction in the reproducibility of the results: The interpretability F1-score with SATORI-Basic demonstrated a standard deviation of 0.0004 across ten runs of the model with different seed values. The addition of the RNN increased this to 0.004, which we would still consider stable.

### 3.2 TF Interactions in Accessible Chromatin in Human Promoters

To test the ability of SATORI to detect cooperation of TFs in real data, we explored the regulatory cooperation among TFs within human promoter regions leveraging DNase I hypersensitivity data (DHSs) collected from 164 immortalized cell lines and tissues [2] as in our earlier work [17]. The task is the prediction of the presence or absence of a DHS across each of the 164 cell lines, leading to a multi-label classification problem. The dataset consists of 20,613 examples across 164 cell types and each sequence is 600bp in length. We partitioned the dataset into training, testing, and validation sets at an approximate ratio of 80%, 10%, and 10%, respectively, while ensuring entire chromosomes were allocated to each set. Specifically, the training set includes chromosomes 1-14 and 18-19, the validation set includes chromosomes 17, 20-21, and the test set contains the remaining chromosomes. To compare the performance of SATORI with trainable CNN filters against pre-loaded CNN weights from clustered PSSMs, as described elsewhere [20], we clustered 733 human transcription factors motifs into 505 groups. The corresponding PSSMs for these clusters were loaded as weights in the first CNN layer. During interaction inference, we followed a similar approach to TIANA [20], where sequences with a filter score greater than half the maximum PSSM score for that filter were considered for further analysis. The mean classification AUC-ROC achieved for the basic and deep SATORI models without entropy were 0.82 and 0.84, respectively. The addition of the entropy term resulted in a slight decrease in classification accuracy, yielding an AUC of 0.79 for the basic model, and 0.81 for the deep model. Models with pre-loaded CNN weights achieved the AUC-ROC of 0.81 with entropy for both basic and deep architectures. However, the entropy term was crucial for detecting interactions: the addition of the entropy term increased the number of interactions detected for both architectures (see Fig. 5). The use of pre-loaded filters provided a further big increase in the number of detected interactions; in this dataset deep and shallow models had similar numbers of detected interactions, as did the attention-attribution and attention-raw models.

**Figure 5:**
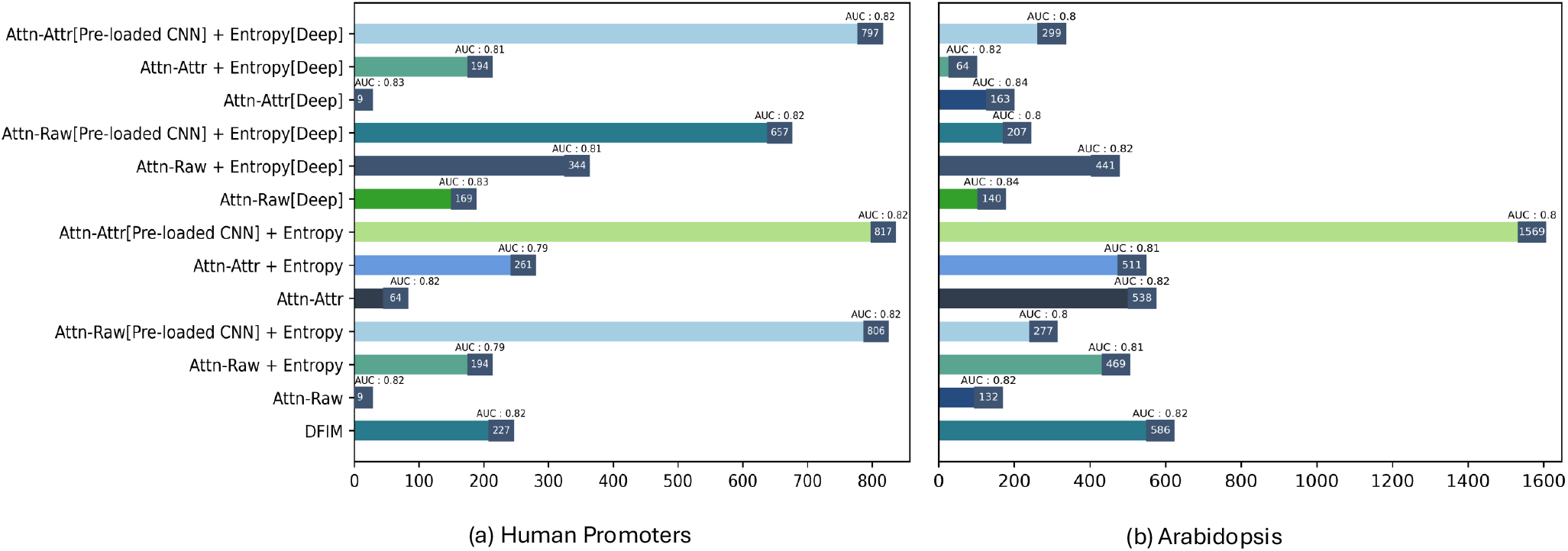
Number of interactions inferred by SATORI basic and deep architectures with and without entropy loss, compared against DFIM on the Human Promoter dataset (a) and accessible chromatin in Arabidopsis (b).

Since the co-occurrence of TF binding can be the result of physical interactions between the TFs, we searched these interactions against the HIPPIE protein-protein interaction database [22]. The interaction between SP7 and KDM2B discovered by all of the methods. Supplementary Table S4 provides the number of matches in the HIPPIE database for all experiments, and we observe that the pre-loaded CNN approach provided the largest number of matches. Tables S5-S16 in the supplementary material provide the list of matches identified. For both real-data experiments, we were unable to run DFIM on deeper models due to time intensive gradient computations.

### 3.3 TF Interactions in Accessible Chromatin in Arabidopsis

For the next experiment, we predicted chromatin accessibility across 36 Arabidopsis samples gathered from DNase I-Seq and ATAC-Seq studies used in our earlier work [17]. This is also a multi-label classification problem with a total of 80,000 instances, each of length 600bp. We partitioned the dataset into training, validation, and testing subsets in an 80%, 10%, and 10% ratio, respectively. Chromosome 2 was allocated to the validation set, chromosome 4 to the test set, and all other chromosomes were included in the training set. We trained two SATORI models, basic and deep, with and without employing the entropy loss. Similar to human promoters data experiments, we clustered 872 Arabidopsis TFs resulting in 343 clusters, which are loaded as CNN weights. The mean classification AUC achieved by both models without entropy on this dataset were 0.82 and 0.83, respectively. For the models trained with entropy loss, basic and deep SATORI models achieved an AUC of 0.77 and 0.82, respectively. Models with pre-loaded CNN weights achieved an AUC of 0.79 on both basic and deep models when using entropy regularization. In this dataset the addition of the entropy term was not as critical for detecting interactions for most of the experiments, and in some cases we obtained a higher number of interactions for the model without entropy. We believe this is a result of the much larger number of examples available in this dataset, making it easier for SATORI to detect significant interactions from the noisy attention weights, thereby requiring less help from the entropy term (Fig. 5(b)). Interestingly, the basic pre-loaded CNN model trained with entropy recovered the highest number of interactions for Arabidopsis (Fig. 5(b)). This can be attributed to the fact that the pre-loaded PSSMs effectively captured the relevant patterns in the data. Out of 343 filters, 342 were activated in more than 10 sequences within the test set, indicating that nearly all motif-based filters were involved during inference. In contrast, for models with trainable CNN filters, when TomTom is unable to match a filter to any known motif, that filter tends to remain inactive during interaction inference. Moreover, when multiple filters are mapped to the same motif, the resulting interactions can become redundant, further reducing the number of inferred interactions. In both Arabidopsis and the human promoters datasets, the attention-based models produced a much larger number of interactions than DFIM.

## 4 Conclusions

In this work we presented SATORI 2.0 which adds the option of an entropy penalty term and incorporates the option of scoring feature interactions based on attribution scores over the attention matrix as an alternative to scoring based on the raw attention values. We performed a comprehensive evaluation of its performance over realistic simulated datasets and real-world datasets with architectures with varying levels of complexity. The results show that the self-attention layers can effectively and accurately discover regulatory interactions across different architectures, datasets and sequence lengths. We found that the entropy loss helps remove false positives, and in most cases, also increases the method’s recall. These findings support entropy as an effective inductive bias for guiding attention towards improved precision when inferring interactions. We also evaluated SATORI models with pre-loaded CNNs as suggested by [20]. Our findings confirm that pre-loaded CNNs perform well in terms of interaction inference due to reduced dependence on Tomtom accuracy. Entropy also appears to improve the precision of models with pre-loaded CNNs. However, using pre-loaded CNNs comes with the challenge of selecting the right set of transcription factors relevant to the problem. In our experiments we found that using attention attribution scores performed similarly to using raw attention values; the earlier DFIM method did not perform as well, and required significantly higher computation times. In summary, the SATORI 2.0 framework provides a comprehensive tool for interpreting neural network layers and obtaining valuable predictions about regulatory relationships that can be further tested in the laboratory.

## 5 Materials and Methods

### 5.1 Simulated Datasets

To obtain reliable estimates of interpretability accuracy we created simulated datasets with implanted interactions. This addresses the lack of ground truth interactions in real-world datasets. In generating our simulated datasets we aimed to replicate the characteristics of real-world datasets. These datasets are designed to be sufficiently complex to challenge the ability of the network to discover interactions. We have generated binary labeled datasets with varying levels of complexity. In each dataset, the positive class is distinguished from the negative class by the presence of interacting pairs of motifs. All sequences are randomly generated based on the nucleotide frequency of the human genome. For each dataset we randomly selected motifs from the JASPAR database [23] and implanted motif occurrences according to their PWMs. In the positive class, for each sequence three interacting pairs are randomly chosen from the predefined set of interacting pairs. These pairs are embedded into the sequence using its PWM, ensuring PWM scores exceeding half of the maximum PWM score. Motifs from an interacting pair are placed in close proximity, within 8-15 base pairs. In the negative class, one to three motifs are randomly selected from the same pool of motifs. These motifs are uniformly embedded using their PWM, with each motif positioned at least 40 base pairs away from every other motif within the sequence. The generated datasets are balanced, consist of a total of 60,000 instances per dataset and each sequence is 300bp long. Table 1 describes the number of TFs embedded in the three simulated datasets along with the number of interactions.

**Table 1:**
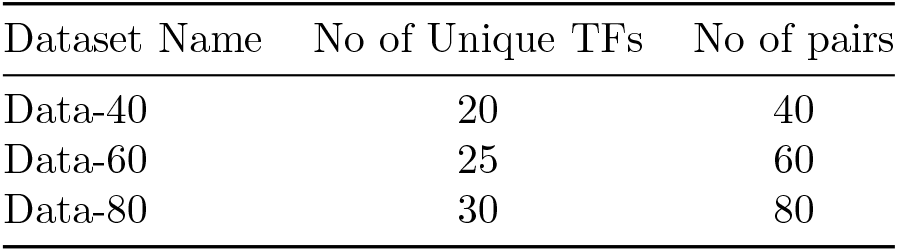
Dataset details. We provide the number of TFs and interacting pairs in the simulated datasets.

| Dataset Name | No of Unique TFs | No of pairs |
| --- | --- | --- |
| Data-40 | 20 | 40 |
| Data-60 | 25 | 60 |
| Data-80 | 30 | 80 |

For the second set of simulated data used to evaluate pre-loaded CNN filters, we used sequences of length 1500bp and motif pairs were drawn from a new set, where Jaspar motifs were clustered and 40 motifs from distinct clusters were selected, and 80 interacting motif pairs were selected. Positive sequences had 3-5 interacting pairs embedded within a window of 8–80bp, while negatives included 1–3 randomly chosen motifs from the pool of unique motifs.

### 5.2 Interpreting Self-Attention Layers with SATORI

SATORI uses a network with a structure that is composed of two or more blocks: an initial block composed of a 1D convolutional feature extraction layer followed by a multi-head self-attention layer; the initial block is followed by additional (optional) deep learning layers and a fully connected output block (see Fig. 1). The attention layer in the primary block serves as a pivotal element for interpretation. Any additional layers serve to provide improved model accuracy, and we demonstrate that this framework can handle models of varying levels of complexity.

The input to the model is a DNA sequence that is represented by its one-hot encoding. The first layer in the network is a one-dimensional convolutional layer that facilitates feature extraction from the DNA sequence. The output of convolution is passed through an activation function. In this work we use the Rectified linear unit function (ReLU) defined as

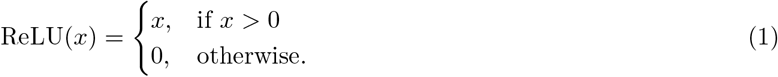

After applying the ReLU function, a max pooling operation is employed to downsample the spatial dimensions of the feature maps. This max-pooling step is optional, and can help in reducing the computation time and memory usage by reducing the length of the sequences that need to be processed by the attention layer, which is the most time and memory consuming aspect of the model. In our experiments we used a very small value for the pooling parameter (4 or less), to ensure that the attention layer receives accurate spatial information about the sequence. The output of the max-pooling operation is fed into the multi-head self-attention layer, which captures relationships between elements within the sequence. This allows the model to weigh the importance of different elements, enabling it to focus on relevant information. Self-attention is computed using three matrices known as query, key, and value defined as:

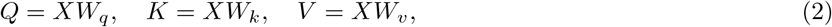

where *X* is the max-pooled output of first CNN layer, *Wq, Wk* and *Wv* are learnable weight matrices used to transform the input vector *X* into the query (*Q*), key (*K*), and value (*V*) matrices. By computing the dot product between the query and key vectors of specific positions in the sequence, we can assess the relationship between the selected positions. We then compute dot product attention as follows:

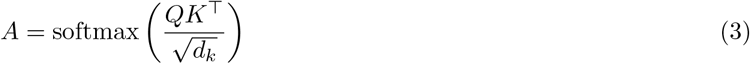

where, *d*_*k*_ is the embedding dimension of *K, Q*, and *V*. The softmax function is used to transform the dot products into probabilities arranged in a row-wise manner. This allows us to interpret the output of the attention layer as the likelihood of a relation between different parts of the input.

The output from the softmax function is employed as weights to compute the value matrix *V* to generate the output of a given attention head:

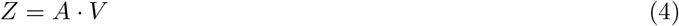

Multi-head attention enables the model to simultaneously attend to different aspects of the input from diverse subspaces across various positions. Subsequently, the outputs of the attention heads are concatenated to produce the final output of the attention layer.

We use a similar process as described by Ullah et al. [17] to discover filter-filter interactions, and infer motif interactions from those filter interactions. To infer interactions we first combine the values of the attention heads by computing the maximum value across heads. Instead of employing a fixed threshold value as was used in the original SATORI, we utilized the mean value of attention scores as a cutoff threshold for selecting interacting attention values. Following this, the statistical significance of the attention values is evaluated using the non-parametric Mann-Whitney U Test [24] relative to the background attention scores computed over the negative examples for binary classification or shuffled sequences in the context of multi-label classification. This process is illustrated in Fig. 1.

We leveraged the weights of the initial CNN layer and attention layer to map filters to putative TFs using an established protocol [2]. Each substring yielding an activation score exceeding 0.65 of the maximum score of the filter is used the final computation of the position weight matrix (PWM) for that filter. The resulting PWMs are then compared against the known motif databases (JASPAR [23] (simulated datasets), Human CISBP [25] (for the human promoters dataset) and Arabidopsis DAP [26]) using the TomTom tool [27], and the top matches are assigned to each filter.

#### A deeper architecture

Our baseline architecture (basic SATORI) directly feeds the output of the attention layer to a fully connected layer. In order to demonstrate the effectiveness of SATORI with more complex architectures we explored the addition of an additional block of convolutional layers whose structure was inspired by the SEI model [28]. The basic component of this block is a residual dual linear and non-linear CNN sub-block shown in Fig. 6. The linear pathway enables rapid training, while the nonlinear pathway is intended to enhance its representational power. Furthermore, a residual connection is established between the stacked linear and nonlinear blocks, thereby permitting computation to traverse either the linear or nonlinear pathway. The last sub-block uses dilated CNN layers augmented with residual connections, to capture longer-range relationships across the sequence. Subsequently, the resulting feature maps are averaged along the sequence dimension prior to processing through a fully connected layer and final output layer.

**Figure 6:**
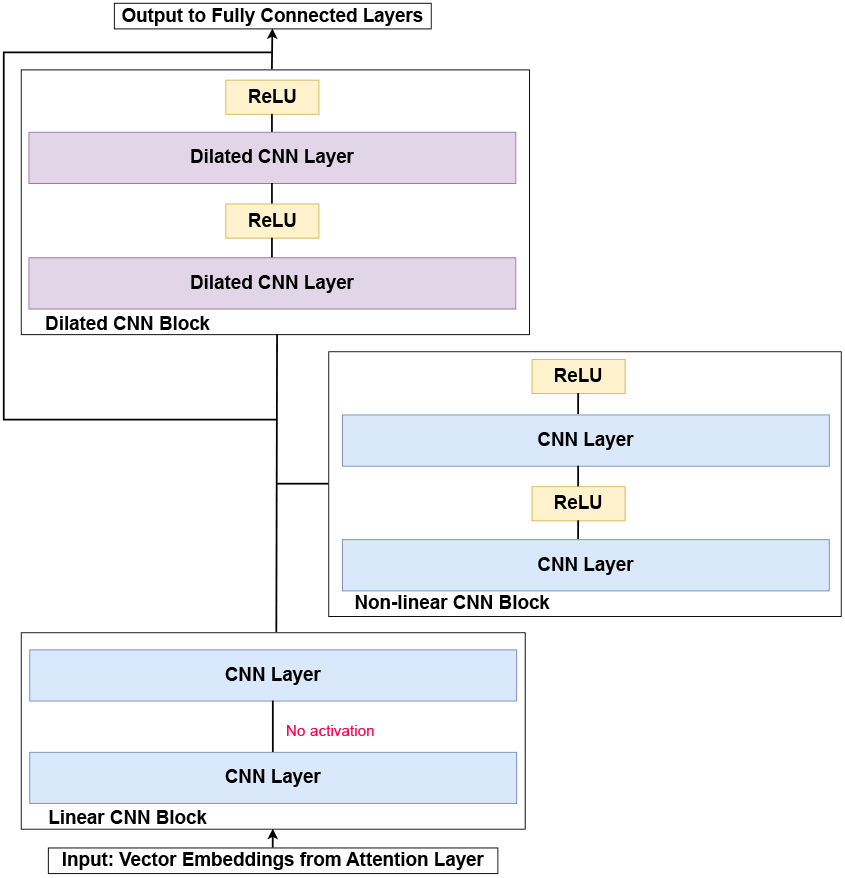
Additional layers included in the SATORI deep architecture after the self-attention layer.

#### Improving network interpretability with an entropy loss term

In order to improve the interpretability of the attention values we incorporated an entropy term in our loss function in addition to the classification loss. The entropy loss is calculated from the attention scores to promote sparsity of the attention matrix. The resulting loss function is defined as follows:

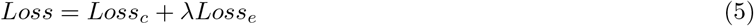

Here, *Loss*_*c*_ is a classification loss (e.g. binary cross entropy) and *Loss*_*e*_ is the entropy term defined as:

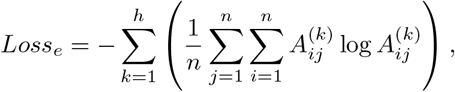

where *n* is number of rows or columns in the attention head *A*^(*k*)^, and *h* is number of heads. The parameter *λ* in Equation 5 is the regularization parameter for the entropy loss and is selected as part of our hyperparameter search.

#### Attention Attribution Scores

To compare the interpretation performance of raw attention scores with attributed attention scores, as used by [19], we incorporated their approach into the SATORI framework. Attention attribution scores are computed by calculating the gradients of the target output with respect to self-attention scores. This method was first proposed by [29] to interpret the information learned by a transformer model. The attribution scores are approximated from a zero-baseline, where no filters attend to each other, to the actual input by gradually increasing the attention scores’ weight. This can be expressed via the Riemann approximation of the integration [30] as follows:

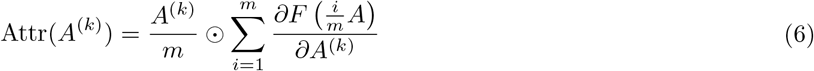

where ⊙ is element-wise multiplication, *A*^(*k*)^ describes a single attention head among a set of *h* attention heads. 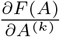 is the gradient of the model *F* (·) with respect to *A*^(*k*)^; *m* is the number of approximation steps, which is set to 20. To identify significant interactions, we process the attention attribution scores in a similar manner to the way SATORI processes raw attention scores. When using the entropy loss with the deep architecture we incorporated residual connections from the output of the feature extraction CNN layer in the initial block to the output of the multihead attention layer before applying additional convolution layers. This is done to maintain the accuracy of the model, which can be impacted by the increased sparsity induced by the entropy loss term.

### 5.3 Model Selection and Training

We used two primary architectures in our experiments: a basic model, which is a two-layered architecture comprising a CNN layer, self-attention layer, and the more complex model which includes additional CNN layers. For model selection for the basic architecture, we employed a randomized grid search algorithm using the predefined search space shown in Table S1 in the supplementary material. This algorithm tunes multiple hyperparameters including the number of CNN filters, CNN filter size, size of the attention head, number of attention heads, and the size of the linear layer after the multi-head attention layer. In addition to these, the optimizer, learning rate and weight decay are also determined through hyperparameter search. An instance of the simulated dataset is used for hyperparameter selection based on its AUC on the validation set. Details of selected models for training are provided in Table S2 of the supplementary material. To obtain an optimal value for the regularization parameter *λ* of the entropy loss, we used a values on a logarithmic scale ranging from 10*−*5 to 10*−*2. We selected the entropy value corresponding to the best interaction performance (F1-score) on the dataset used for model selection for further experimentation. Datasets that are used for model and hyperparameter selection are later discarded, and new instances are utilized to evaluate the performance of the final selected model. For the real datasets we used architecture settings similar to our previous work [17]; the deep architecture used the same hyperparameters for the initial block; details of the SEI-inspired additional block are given in Table S3 in supplementary material. The simulated datasets were divided into train, validation and test sets with 80%, 10% and 10% ratios, respectively. For each dataset size we generated three dataset instances. Subsequently, we trained a model three times using different seed values on each instance of the dataset, and the best-performing model, determined by validation AUC was selected. The final results for each dataset type are reported by averaging over the three dataset instances. Model accuracy was measured using area under the ROC curve (AUC) and interpretation accuracy of the predicted interactions was measured using precision, recall, and F1-score. All methods were implemented using PyTorch [31] and all the experiments were conducted on a Linux workstation with a 12GB TITAN V GPU. For DFIM implementation, we have used the implementation provided in the original SATORI code base [17].

## 6 Data Availability

The source code and datasets are available at https://github.com/sairajab/satoriv2. A snapshot of the repository is available at https://doi.org/10.5281/zenodo.13362826.

## References

[1] Jian Zhou and Olga G Troyanskaya. Predicting effects of noncoding variants with deep learning–based sequence model. Nature methods, 12(10):931–934, 2015.

[2] David R Kelley, Jasper Snoek, and John L Rinn. Basset: learning the regulatory code of the accessible genome with deep convolutional neural networks. Genome research, 26(7):990–999, 2016.

[3] Žiga Avsec, Vikram Agarwal, Daniel Visentin, Joseph R Ledsam, Agnieszka Grabska-Barwinska, Kyle R Taylor, Yannis Assael, John Jumper, Pushmeet Kohli, and David R Kelley. Effective gene expression prediction from sequence by integrating long-range interactions. Nature methods, 18(10):1196–1203, 2021.

[4] Ritambhara Singh, Jack Lanchantin, Gabriel Robins, and Yanjun Qi. Deepchrome: deep-learning for predicting gene expression from histone modifications. Bioinformatics, 32(17):i639–i648, 2016.

[5] Alireza Karbalayghareh, Merve Sahin, and Christina S Leslie. Chromatin interaction–aware gene regulatory modeling with graph attention networks. Genome Research, 32(5):930–944, 2022.

[6] Somaye Hashemifar, Behnam Neyshabur, Aly A Khan, and Jinbo Xu. Predicting protein–protein interactions through sequence-based deep learning. Bioinformatics, 34(17):i802–i810, 2018.

[7] John Jumper, Richard Evans, Alexander Pritzel, Tim Green, Michael Figurnov, Olaf Ronneberger, Kathryn Tunyasuvunakool, Russ Bates, Augustin Z?ídek, Anna Potapenko, et al. Highly accurate protein structure prediction with AlphaFold. Nature, 596(7873):583–589, 2021.

[8] Babak Alipanahi, Andrew Delong, Matthew T Weirauch, and Brendan J Frey. Predicting the sequence specificities of DNA-and RNA-binding proteins by deep learning. Nature biotechnology, 33(8):831–838, 2015.

[9] Mukund Sundararajan, Ankur Taly, and Qiqi Yan. Axiomatic attribution for deep networks. In International conference on machine learning, pages 3319–3328. PMLR, 2017.

[10] Anupama Jha, Joseph K. Aicher, Matthew R. Gazzara, Deependra Singh, and Yoseph Barash. Enhanced integrated gradients: improving interpretability of deep learning models using splicing codes as a case study. Genome biology, 21:1–22, 2020.

[11] Wyeth W Wasserman and James W Fickett. Identification of regulatory regions which confer muscle-specific gene expression. Journal of molecular biology, 278(1):167–181, 1998.

[12] Satyanarayan Rao, Kami Ahmad, and Srinivas Ramachandran. Cooperative binding of transcription factors is a hallmark of active enhancers. bioRxiv, pages 2020–08, 2020.

[13] Priya Sudarsanam, Yitzhak Pilpel, and George M Church. Genome-wide co-occurrence of promoter elements reveals a cis-regulatory cassette of rRNA transcription motifs in saccharomyces cerevisiae. Genome research, 12(11):1723–1731, 2002.

[14] Ignacio L Ibarra, Nele M Hollmann, Bernd Klaus, Sandra Augsten, Britta Velten, Janosch Hennig, and Judith B Zaugg. Mechanistic insights into transcription factor cooperativity and its impact on protein-phenotype interactions. Nature communications, 11(1):124, 2020.

[15] Peyton Greenside, Tyler Shimko, Polly Fordyce, and Anshul Kundaje. Discovering epistatic feature interactions from neural network models of regulatory DNA sequences. Bioinformatics, 34(17):i629–i637, 2018.

[16] Avanti Shrikumar, Peyton Greenside, and Anshul Kundaje. Learning important features through propagating activation differences. In International conference on machine learning, pages 3145–3153. PMLR, 2017.

[17] Fahad Ullah and Asa Ben-Hur. A self-attention model for inferring cooperativity between regulatory features. Nucleic acids research, 49(13):e77–e77, 2021.

[18] Zhenhao Zhang, Fan Feng, Yiyang Qiu, and Jie Liu. A generalizable framework to comprehensively predict epigenome, chromatin organization, and transcriptome. Nucleic Acids Research, 51(12):5931–5947, 2023.

[19] Anil Prakash and Moinak Banerjee. An interpretable block-attention network for identifying regulatory feature interactions. Briefings in Bioinformatics, 24(4):bbad250, 2023.

[20] Rick Z Li, Claudia Z Han, and Christopher K Glass. TIANA: transcription factors cooperativity inference analysis with neural attention. BMC bioinformatics, 25(1):274, 2024.

[21] Zhenhao Zhang, Fan Feng, and Jie Liu. Characterizing collaborative transcription regulation with a graph-based deep learning approach. PLOS Computational Biology, 18(6):e1010162, 2022.

[22] Gregorio Alanis-Lobato, Miguel A Andrade-Navarro, and Martin H Schaefer. HIPPIE v2. 0: enhancing meaningfulness and reliability of protein–protein interaction networks. Nucleic acids research, page gkw985, 2016.

[23] Oriol Fornes, Jaime A Castro-Mondragon, Aziz Khan, Robin Van der Lee, Xi Zhang, Phillip A Richmond, Bhavi P Modi, Solenne Correard, Marius Gheorghe, Damir Barana?síc, et al. JASPAR 2020: update of the open-access database of transcription factor binding profiles. Nucleic acids research, 48(D1):D87–D92, 2020.

[24] Henry B Mann and Donald R Whitney. On a test of whether one of two random variables is stochastically larger than the other. The annals of mathematical statistics, pages 50–60, 1947.

[25] Matthew T Weirauch, Ally Yang, Mihai Albu, Atina G Cote, Alejandro Montenegro-Montero, Philipp Drewe, Hamed S Najafabadi, Samuel A Lambert, Ishminder Mann, Kate Cook, et al. Determination and inference of eukaryotic transcription factor sequence specificity. Cell, 158(6):1431–1443, 2014.

[26] Ronan C O’Malley, Shao-shan Carol Huang, Liang Song, Mathew G Lewsey, Anna Bartlett, Joseph R Nery, Mary Galli, Andrea Gallavotti, and Joseph R Ecker. Cistrome and epicistrome features shape the regulatory DNA landscape. Cell, 165(5):1280–1292, 2016.

[27] Shobhit Gupta, John A Stamatoyannopoulos, Timothy L Bailey, and William Stafford Noble. Quantifying similarity between motifs. Genome biology, 8:1–9, 2007.

[28] Kathleen M Chen, Aaron K Wong, Olga G Troyanskaya, and Jian Zhou. A sequence-based global map of regulatory activity for deciphering human genetics. Nature genetics, 54(7):940–949, 2022.

[29] Yaru Hao, Li Dong, Furu Wei, and Ke Xu. Self-attention attribution: Interpreting information interactions inside transformer. In Proceedings of the AAAI Conference on Artificial Intelligence, volume 35, pages 12963–12971, 2021.

[30] Mukund Sundararajan, Ankur Taly, and Qiqi Yan. Axiomatic attribution for deep networks. In Doina Precup and Yee Whye Teh, editors, Proceedings of the 34th International Conference on Machine Learning, volume 70 of Proceedings of Machine Learning Research, pages 3319–3328. PMLR, 06–11 Aug 2017.

[31] Adam Paszke, Sam Gross, Francisco Massa, Adam Lerer, James Bradbury, Gregory Chanan, Trevor Killeen, Zeming Lin, Natalia Gimelshein, Luca Antiga, et al. PyTorch: An imperative style, high-performance deep learning library. Advances in neural information processing systems, 32, 2019.

