## Supplementary material for "A Comprehensive Evaluation of Self Attention for Detecting Regulatory Feature Interactions"

### SUPPLEMENTARY FIGURES

Seq 1

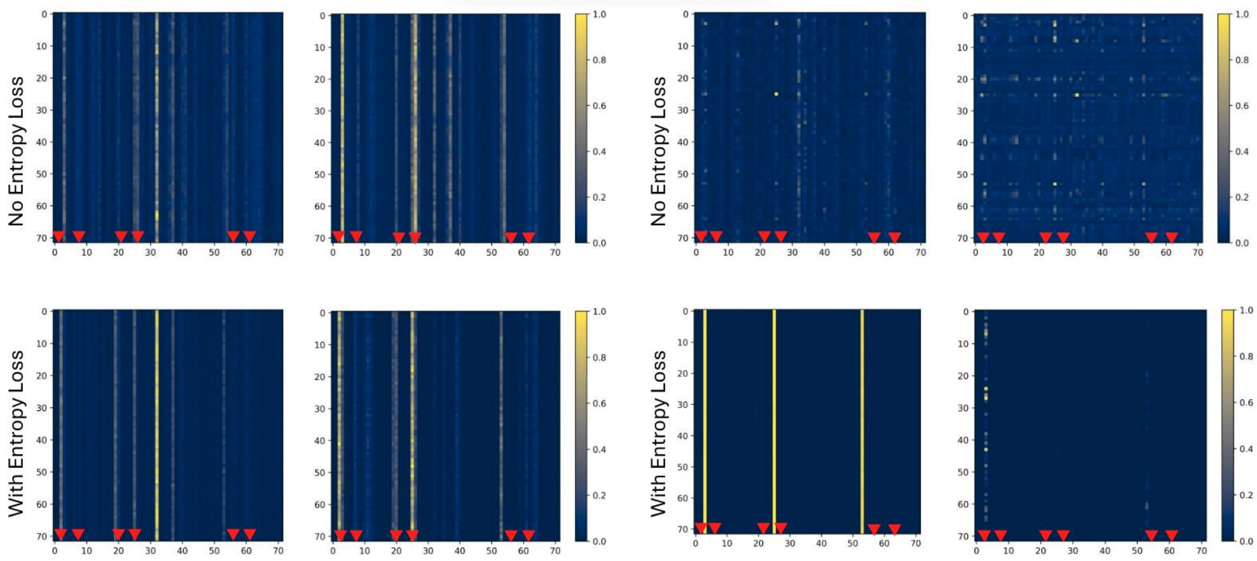


(a)Raw Attention scores (Basic) (b)Attention Attribution scores (Basic) (c)Raw Attention scores (Deep) (d)Attention Attribution scores (Deep)

Seq 2

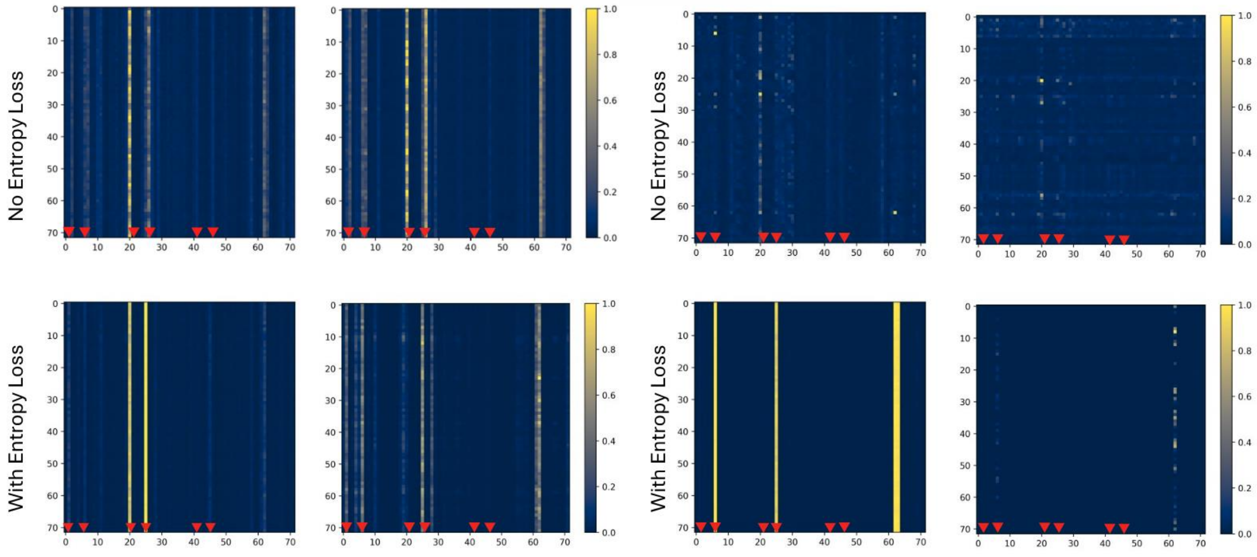


(a)Raw Attention scores (Basic) (b)Attention Attribution scores (Basic) (c)Raw Attention scores (Deep) (d)Attention Attribution scores (Deep)

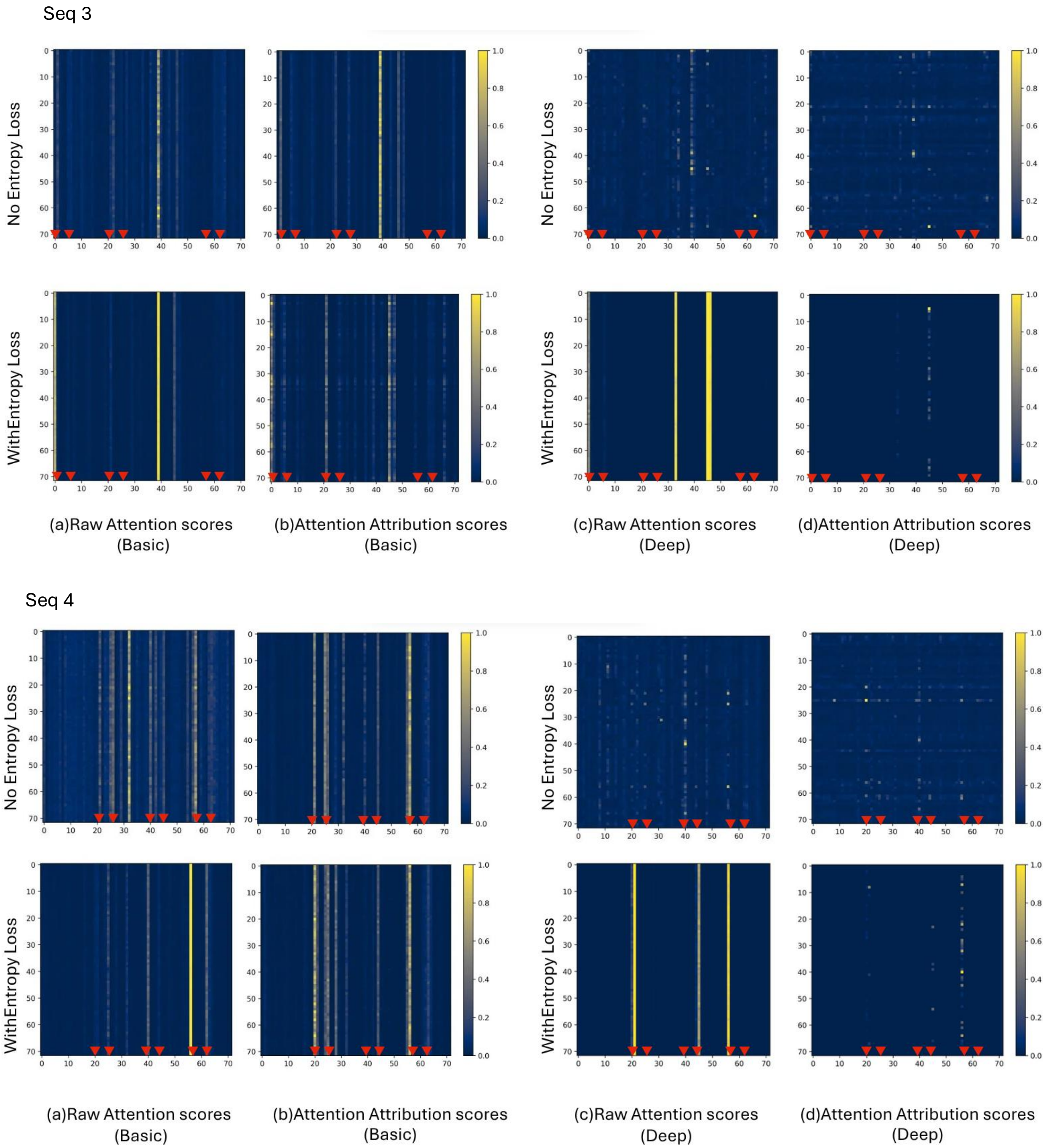


**Fig. S2.** Comparison of heatmaps showing raw attention scores (a, c) and attention attribution scores (b, d), max-pooled over attention heads, for four true positive test sequences from Data-40 across four different models. The top row represents models trained without entropy loss, while the bottom row shows models trained with entropy loss. (a) and (b) depict the Basic architecture, whereas (c) and (d) illustrate the Deep architecture. In all heatmaps, both axes represent positions within the sequence after max-pooling. Each position on the Y-axis corresponds to attention probabilities with respect to each position on the X-axis. Vertical lines on the X-axis indicate embedded motif positions. This visualization enables comparison of attention patterns across different model architectures, training strategies, and score types, highlighting the impact of entropy loss and architecture depth on attention distribution. Red arrows show the positions where motifs are embedded, adjusted for pooling window.

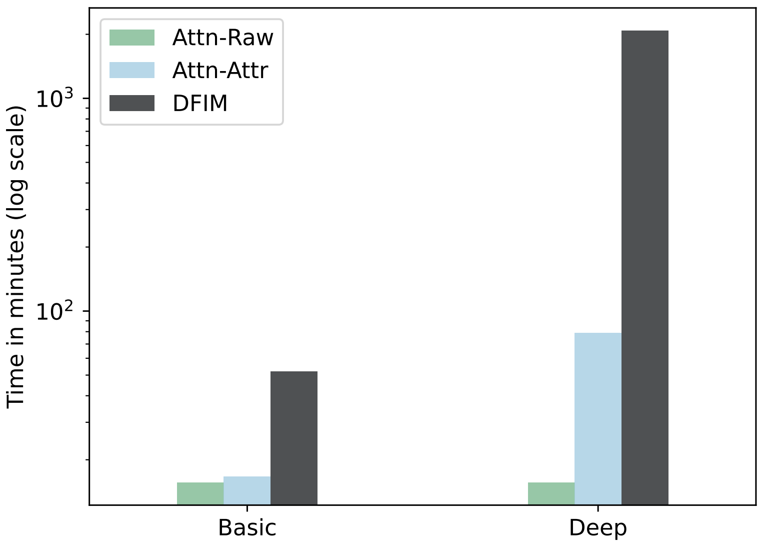


**Fig. S3.** Average runtime for SATORI with raw attention scores, attention attribution scores, and DFIM to estimate interactions for Data-40.

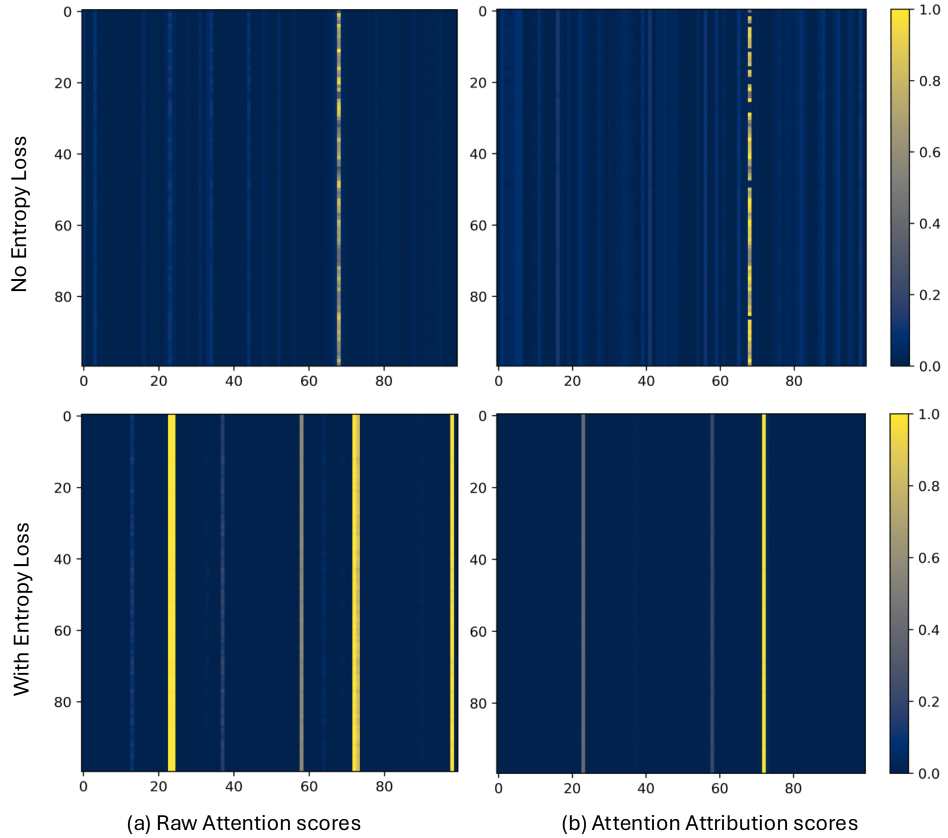


**Fig. S4.** Heatmaps comparing raw attention scores (a) and attention attribution scores (b) for a true positive test sequence from the Human Promoters dataset using the Basic architecture. (a) and (b) demonstrate similar patterns. The top row shows less sparsity in the raw attention scores and attention attribution scores for a model trained without entropy loss compared to the bottom row, which reveals a bimodal representation of the attention layer when trained with entropy loss.

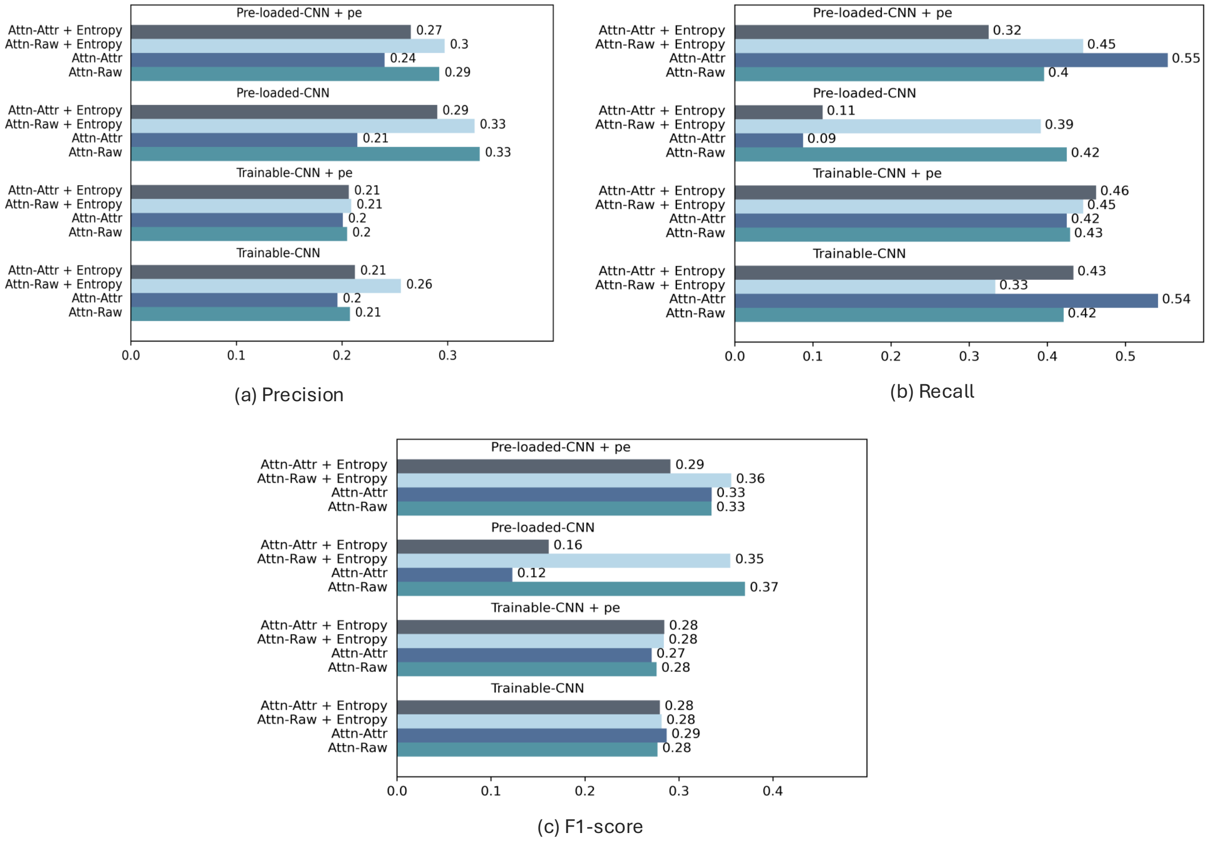


**Fig. S5.** Comparative performance of Basic-SATORI model on simulated datasets with longer sequences (1500 bp) containing 80 motif pairs. We report interpretation performance as: Precision (a), Recall (b) and F1-score (c). Results are shown for eight model variants: Basic-SATORI with trainable or pre-loaded CNN filters, each trained with or without entropy loss, and with or without positional encoding. Performance metrics are averaged over three dataset instances and reported using both raw attention scores (Attn-Raw) and attention attribution scores (Attn-Attr).

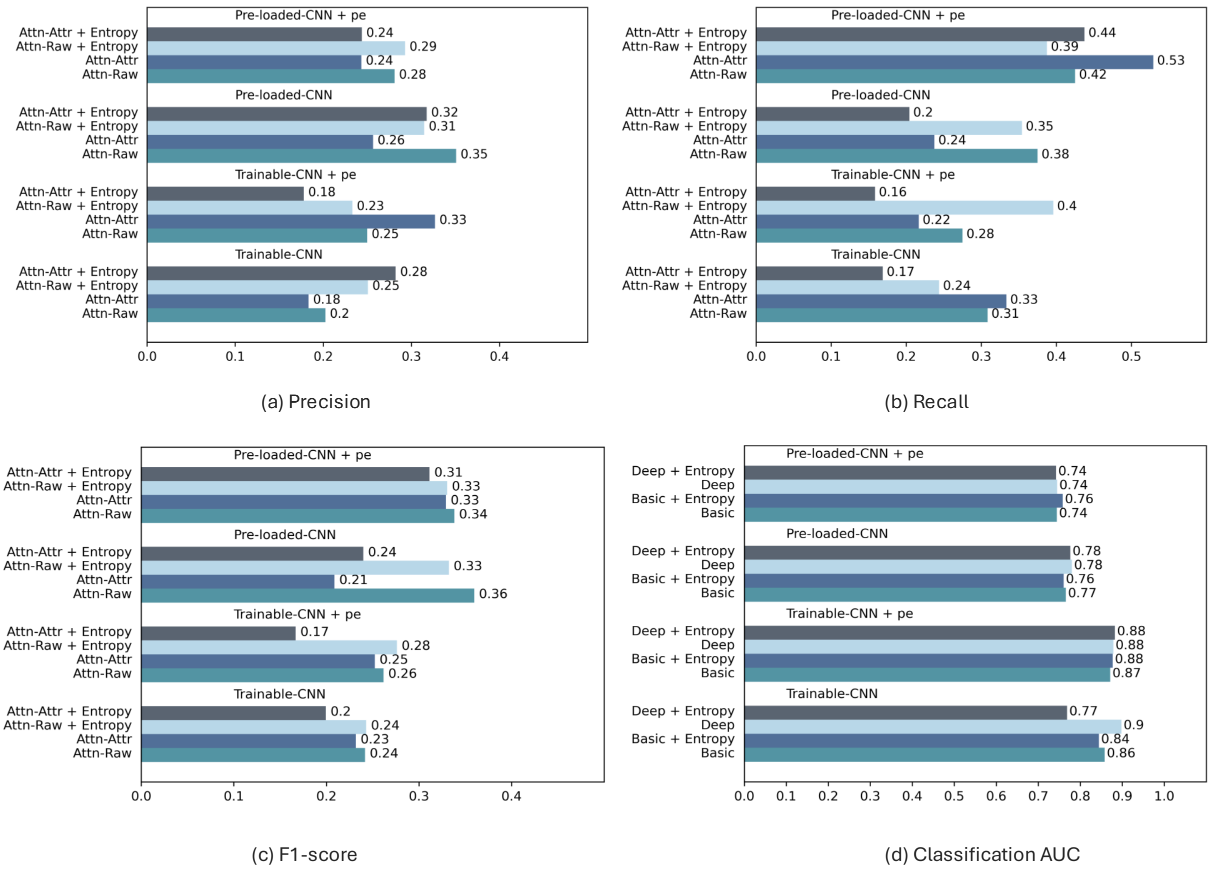


(a) Precision

(b) Recall

(c) F1-score

(d) Classification AUC

**Fig. S6.** Comparative performance of Deep-SATORI model on simulated datasets with longer sequences (1500 bp) containing 80 motif pairs. We report interpretation perfromance as: Precision (a), Recall (b) F1-score (c) and classification AUC (d). Results are shown for eight model variants: Deep-SATORI with trainable or pre-loaded CNN filters, each trained with or without entropy loss, and with or without positional encoding. Performance metrics are averaged over three dataset instances and reported using both raw attention scores (Attn-Raw) and attention attribution scores (Attn-Attr).

**Positional Encoding:** We used positional encoding to inject position information into the output of the CNN Layer as used in [1]. The encoding uses sine and cosine functions of different frequencies, as originally proposed in [2].

### SUPPLEMENTARY TABLES

**Table S1.** Basic SATORI architecture's hyper-parameters, search space and sampling method

| Hyper-parameters | Search space | Sampling |
|---|---|---|
| No. of CNN filters | [200, 300, 400] | Uniform |
| CNN Filter Size | [13, 15, 17, 19] | Uniform |
| No. of heads | [2, 4, 8] | Uniform |
| Head Size | [32, 64, 128] | Uniform |
| FC Layer Size | [100, 200, 256,512] | Uniform |
| No of FC layers | [1,2,4] | Uniform |
| Layer Normalization | [Yes, No] | Uniform |
| Optimizer | [SGD, Adagrad, Adam] | Uniform |
| Learning Rate | [0.0001,0.1] | Log uniform |
| Momentum (SGD) | [0.95,0.99] | Sqrt uniform |
| Weight Decay | $[10^{-6}, 10^{-1}]$ | Log uniform |
| Batch Size | [32, 64, 128, 256] | Uniform |
| No of epochs | [30, 40, 50, 60] | Uniform |
| Intialization Seed | [0,1,2] | Evaluate all |

**Table S2.** Architecture specifications of basic and deep SATORI models for simulated and real datasets

| **Architecture specs** | | **Basic (Simulated)** | **Deep (Simulated)** | **Basic (Real dataset)** | **Deep (Real dataset)** |
|---|---|---|---|---|---|
| Conv1d | # of filters | 200 | 256 | 200 | 256 |
| | Filter size | 13 | 13 | 13 | 13 |
| | Padding | 0 | 0 | 6 | 6 |
| Batch Normalization & ReLU() | Same for all | | | | |
| MaxPool1d | Filter size | 4 | 4 | 6 | 6 |
| | Padding | 0 | 0 | 6 | 6 |
| Dropout | 0.2 | | | | |
| Multihead Attention | Head size | 64 | 64 | 32 | 32 |
| | # of heads | 4 | 4 | 8 | 8 |
| LayerNorm | | Yes | No | No | No |
| Linear 1 | Size | 512 | 512 | 1024 | 1024 |
| ReLU() & Dropout | 0.2 | | | | |
| Linear 2 | Size | 100 | **Supplementary Layers Block** | - | **Supplementary Layers Block** |
| ReLU () & Dropout | | 0.2 | | - | |
| Readout | Normalize | | | | |
| Linear 3 | Size | **2** | | **No of classes** | |

**Table S3.** SEI-inspired Supplementary Layers Block

| **Layer** | **# of filters** | **Filter size** | **Padding** | **Dilation** |
|---|---|---|---|---|
| Conv1d | 480 | 9 | 4 | - |
| Conv1d | 480 | 9 | 4 | - |
| Conv1d | 480 | 9 | 4 | - |
| ReLU() | | | | |
| Conv1d | 480 | 9 | 4 | - |
| ReLU() | | | | |
| Dropout(0.4) | | | | |
| Dconv1d | 480 | 5 | 4 | 2 |
| ReLU() | | | | |
| Dropout (0.4) | | | | |
| Dconv1d | 480 | 5 | 8 | 4 |
| ReLU() | | | | |
| **Output Layer** | | | | |

**Table S4.** Number of matched interactions inferred by different methods with HIPPIE databases

| Methods | Matches in HIPPIE |
|---|---|
| DFIM | 3 |
| Attn-Raw | 1 |
| Attn-Raw + Entropy | 3 |
| Attn-Raw[Pre-loaded CNN] + Entropy | 30 |
| Attn-Attr | 2 |
| Attn-Attr + Entropy | 4 |
| Attn-Attr[Pre-loaded CNN] + Entropy | 27 |
| Attn-Raw[Deep] | 5 |
| Attn-Raw + Entropy[Deep] | 4 |
| Attn-Raw[Pre-loaded CNN] + Entropy[Deep] | 21 |
| Attn-Attr[Deep] | 0 |
| Attn-Attr + Entropy[Deep] | 4 |
| Attn-Attr[Pre-loaded CNN] + Entropy[Deep] | 25 |

**Table S5.** Significant matches with HIPPIE database inferred by SATORI basic using Attn-Raw scores on Human Promoters Dataset

| TF Interaction | TF1 Family | TF2 Family | P-value |
|---|---|---|---|
| SP7⟷KDM2B | C2H2 ZF | CxxC | $3.50 \times 10^{-5}$ |

**Table S6.** Significant matches with HIPPIE database inferred by SATORI basic using Attn-Attr scores on Human Promoters Dataset

| TF Interaction | TF1 Family | TF2 Family | P-value |
|---|---|---|---|
| KDM2B⟷SP7 | CxxC | C2H2 ZF | $9.00 \times 10^{-4}$ |
| ZFY⟷ZFX | C2H2 ZF | C2H2 ZF | $5.00 \times 10^{-5}$ |

**Table S7.** Significant matches with HIPPIE database inferred by SATORI basic using DFIM on Human Promoters Dataset

| TF Interaction | TF1 Family | TF2 Family | P-value |
|---|---|---|---|
| TLX2⟷KDM2B | Homeodomain | CxxC | $8.71 \times 10^{-4}$ |
| KDM2B⟷SP7 | CxxC | C2H2 ZF | $5.56 \times 10^{-4}$ |
| ZFY⟷ZFX | C2H2 ZF | C2H2 ZF | $4.90 \times 10^{-3}$ |

**Table S8.** Significant matches with HIPPIE database inferred by SATORI basic with Entropy Loss using Attn-Raw scores on Human Promoters Dataset

| TF Interaction | TF1 Family | TF2 Family | P-value |
|---|---|---|---|
| SP7⟷KDM2B | C2H2 ZF | CxxC | $8.95 \times 10^{-3}$ |
| SP7⟷SIX4 | C2H2 ZF | Homeodomain | $6.56 \times 10^{-3}$ |
| TLX2⟷KDM2B | Homeodomain | CxxC | $9.24 \times 10^{-3}$ |

**Table S9.** Significant matches with HIPPIE database inferred by SATORI basic with Entropy Loss using Attn-Attr scores on Human Promoters Dataset

| TF Interaction | TF1 Family | TF2 Family | P-value |
|---|---|---|---|
| SP7⟷KDM2B | C2H2 ZF | CxxC | $1.65 \times 10^{-4}$ |
| SP7⟷SIX4 | C2H2 ZF | Homeodomain | $2.99 \times 10^{-2}$ |
| ZFX⟷ZFY | C2H2 ZF | C2H2 ZF | $7.80 \times 10^{-11}$ |
| TLX2⟷KDM2B | Homeodomain | CxxC | $5.01 \times 10^{-3}$ |

**Table S10.** Significant matches with HIPPIE database inferred by SATORI basic with Pre-loaded CNN and Entropy Loss using Attn-Raw scores on Human Promoters Dataset

| TF Interaction | TF1 Family | TF2 Family | P-value |
|---|---|---|---|
| PPARA, RARA, RARB⟷ESR1 | Nuclear receptor | Nuclear receptor | $3.69 \times 10^{-2}$ |
| PPARA, RARA, RARB⟷ESR2, ESRRA | Nuclear receptor | Nuclear receptor | $3.69 \times 10^{-2}$ |
| MECP2⟷PPARG | MBD | Nuclear receptor | $4.13 \times 10^{-3}$ |
| NR0B1⟷ESR1 | Unknown | Nuclear receptor | $4.13 \times 10^{-3}$ |
| NR0B1⟷PPARG | Unknown | Nuclear receptor | $8.76 \times 10^{-3}$ |
| ID2, NPAS2, MLXIP, MYC, MNT⟷TFDP1 | bHLH | DP,E2F | $8.76 \times 10^{-3}$ |
| EPAS1⟷USF2 | bHLH | bHLH | $3.69 \times 10^{-2}$ |
| ERF, ETV1, ELK1, ETV3⟷NR2C2 | Ets | Nuclear receptor | $1.80 \times 10^{-2}$ |
| NR2C1⟷NR2C2 | Nuclear receptor | Nuclear receptor | $3.69 \times 10^{-2}$ |
| SP2, SP1⟷NR2C2 | C2H2 ZF | Nuclear receptor | $3.69 \times 10^{-6}$ |
| SP2, SP1⟷ETS1, SMARCC2 | C2H2 ZF | Ets, Myb/SANT | $9.57 \times 10^{-4}$ |
| SP2, SP1⟷REST | C2H2 ZF | C2H2 ZF | $3.69 \times 10^{-2}$ |
| SP5, KLF12, KLF7⟷NR2C2 | C2H2 ZF | Nuclear receptor | $9.42 \times 10^{-8}$ |
| JUN⟷PURA | bZIP | Unknown | $3.69 \times 10^{-2}$ |
| WT1⟷NR2C2 | C2H2 ZF | Nuclear receptor | $1.32 \times 10^{-5}$ |
| SMAD2⟷ESR2, ESRRA | SMAD | Nuclear receptor | $4.13 \times 10^{-3}$ |
| TGIF1⟷SMAD1 | Homeodomain | SMAD | $3.69 \times 10^{-2}$ |
| SMAD3, SMAD4⟷PPARG | SMAD | Nuclear receptor | $4.13 \times 10^{-3}$ |
| BRCA1, ZBTB33⟷E2F1 | C2H2 ZF, EIN3 | E2F | $1.80 \times 10^{-2}$ |
| RFX4⟷RFX2 | RFX | RFX | $8.76 \times 10^{-3}$ |
| THRA, RXRB⟷NR2C2 | Nuclear receptor | Nuclear receptor | $2.02 \times 10^{-3}$ |
| ESR1⟷SP3 | Nuclear receptor | C2H2 ZF | $8.76 \times 10^{-3}$ |
| PPARG⟷ZBTB3 | Nuclear receptor | C2H2 ZF | $2.29 \times 10^{-4}$ |
| NR2C2⟷MAF, CIC, NRL | Nuclear receptor | Sox, bZIP | $1.32 \times 10^{-5}$ |
| NR2C2⟷SP8, KLF14, KLF16 | Nuclear receptor | C2H2 ZF | $2.29 \times 10^{-4}$ |
| NR2C2⟷PAX8 | Nuclear receptor | Paired box | $3.69 \times 10^{-2}$ |
| TFAP4⟷CTCF | bHLH | C2H2 ZF | $1.80 \times 10^{-2}$ |
| ETS1, SMARCC2⟷KDM2B | Ets, Myb/SANT | CxxC | $5.19 \times 10^{-5}$ |
| ETS1, SMARCC2⟷ETS2 | Ets, Myb/SANT | Ets | $4.73 \times 10^{-4}$ |
| PURA⟷ESR2, ESRRA | Unknown | Nuclear receptor | $8.76 \times 10^{-3}$ |

**Table S11.** Significant matches with HIPPIE database inferred by SATORI basic with Pre-loaded CNN and Entropy Loss using Attn-Attr scores on Human Promoters Dataset

| TF Interaction | TF1 Family | TF2 Family | P-value |
|---|---|---|---|
| MECP2⟷PPARG | MBD | Nuclear receptor | $4.69 \times 10^{-3}$ |
| NR0B1⟷ESR1 | Unknown | Nuclear receptor | $1.01 \times 10^{-2}$ |
| NR0B1⟷PPARG | Unknown | Nuclear receptor | $2.08 \times 10^{-2}$ |
| ID2, NPAS2, MLXIP, MYC, MNT⟷TFDP1 | bHLH | DP,E2F | $1.01 \times 10^{-2}$ |
| EPAS1⟷USF2 | bHLH | bHLH | $4.33 \times 10^{-2}$ |
| ERF, ETV1, ELK1, ETV3⟷NR2C2 | Ets | Nuclear receptor | $2.08 \times 10^{-2}$ |
| NR2C1⟷NR2C2 | Nuclear receptor | Nuclear receptor | $4.33 \times 10^{-2}$ |
| SP2, SP1⟷NR2C2 | C2H2 ZF | Nuclear receptor | $4.82 \times 10^{-6}$ |
| SP2, SP1⟷ETS1, SMARCC2 | C2H2 ZF | Ets, Myb/SANT | $1.13 \times 10^{-3}$ |
| SP2, SP1⟷REST | C2H2 ZF | C2H2 ZF | $4.33 \times 10^{-2}$ |
| NFIA, NFIB, NFIX⟷SP8, KLF14, KLF16 | SMAD | C2H2 ZF | $4.33 \times 10^{-2}$ |
| SP5, KLF12, KLF7⟷NR2C2 | C2H2 ZF | Nuclear receptor | $1.17 \times 10^{-7}$ |
| JUN⟷PURA | bZIP | Unknown | $1.01 \times 10^{-2}$ |
| WT1⟷NR2C2 | C2H2 ZF | Nuclear receptor | $3.09 \times 10^{-5}$ |
| SMAD2⟷ESR2, ESRRA | SMAD | Nuclear receptor | $4.69 \times 10^{-3}$ |
| TGIF1⟷SMAD1 | Homeodomain | SMAD | $4.33 \times 10^{-2}$ |
| SMAD3, SMAD4⟷PPARG | SMAD | Nuclear receptor | $1.01 \times 10^{-2}$ |
| RFX4⟷RFX2 | RFX | RFX | $1.01 \times 10^{-2}$ |
| THRA, RXRB⟷NR2C2 | Nuclear receptor | Nuclear receptor | $2.33 \times 10^{-3}$ |
| PPARG⟷ZBTB3 | Nuclear receptor | C2H2 ZF | $5.54 \times 10^{-4}$ |
| NR2C2⟷MAF, CIC, NRL | Nuclear receptor | Sox, bZIP | $3.09 \times 10^{-5}$ |
| NR2C2⟷SP8, KLF14, KLF16 | Nuclear receptor | C2H2 ZF | $2.79 \times 10^{-4}$ |
| NR2C2⟷PAX8 | Nuclear receptor | Paired box | $4.33 \times 10^{-2}$ |
| TFAP4⟷CTCF | bHLH | C2H2 ZF | $2.08 \times 10^{-2}$ |
| ETS1, SMARCC2⟷KDM2B | Ets, Myb/SANT | CxxC | $6.33 \times 10^{-5}$ |
| ETS1, SMARCC2⟷ETS2 | Ets, Myb/SANT | Ets | $5.54 \times 10^{-4}$ |
| PURA⟷ESR2, ESRRA | Unknown | Nuclear receptor | $4.69 \times 10^{-3}$ |

**Table S12.** Significant matches with HIPPIE database inferred by SATORI deep using Attn-Raw scores on Human Promoters Dataset

| TF Interactions | TF1 Family | TF2 Family | P-value |
|---|---|---|---|
| TLX2⟷KDM2B | Homeodomain | CxxC | $6.13 \times 10^{-3}$ |
| SP7⟷KDM2B | C2H2 ZF | CxxC | $4.58 \times 10^{-4}$ |
| SP7⟷SIX4 | C2H2 ZF | Homeodomain | $5.79 \times 10^{-3}$ |
| KDM2B⟷ESR1 | CxxC | Nuclear receptor | $6.52 \times 10^{-3}$ |
| ESR1⟷ZHX1 | Nuclear receptor | Homeodomain | $2.06 \times 10^{-3}$ |

**Table S13.** Significant matches with HIPPIE database inferred by SATORI deep with Entropy Loss using Attn-Raw scores on Human Promoters Dataset

| TF Interactions | TF1 Family | TF2 Family | P-value |
|---|---|---|---|
| SP7⟷FUBP1 | C2H2 ZF | Unknown | $3.77 \times 10^{-3}$ |
| SP7⟷KDM2B | C2H2 ZF | CxxC | $7.29 \times 10^{-3}$ |
| FOXP1⟷SP7 | Forkhead | C2H2 ZF | $6.60 \times 10^{-4}$ |
| ZFY⟷ZFX | C2H2 ZF | C2H2 ZF | $2.75 \times 10^{-4}$ |

**Table S14.** Significant matches with HIPPIE database inferred by SATORI deep with Entropy Loss using Attn-Attr scores on Human Promoters Dataset

| TF Interactions | TF1 Family | TF2 Family | Adjusted P-value |
|---|---|---|---|
| FOXP1⟷SP7 | Forkhead | C2H2 ZF | $3.80 \times 10^{-3}$ |
| SP7⟷KDM2B | C2H2 ZF | CxxC | $2.28 \times 10^{-5}$ |
| SP7⟷FUBP1 | C2H2 ZF | Unknown | $4.10 \times 10^{-3}$ |
| ZFY⟷ZFX | C2H2 ZF | C2H2 ZF | $1.35 \times 10^{-6}$ |

**Table S15.** Significant matches with HIPPIE database inferred by SATORI deep with Pre-loaded CNN and Entropy Loss using Attn-Raw scores on Human Promoters Dataset

| TF Interaction | TF1 Family | TF2 Family | P-value |
|---|---|---|---|
| MBD2⟷ESR1 | MBD | Nuclear receptor | $4.27 \times 10^{-3}$ |
| NR0B1⟷ESR1 | Unknown | Nuclear receptor | $4.27 \times 10^{-3}$ |
| SP2, SP1⟷ESR1 | C2H2 ZF | Nuclear receptor | $4.27 \times 10^{-3}$ |
| SP2, SP1⟷ETS1, SMARCC2 | C2H2 ZF | Ets, Myb/SANT | $1.05 \times 10^{-3}$ |
| NFIA, NFIB, NFIX⟷SMAD2 | SMAD | SMAD | $3.61 \times 10^{-2}$ |
| JUN⟷PURA | bZIP | Unknown | $1.77 \times 10^{-2}$ |
| WT1⟷NR2C2 | C2H2 ZF | Nuclear receptor | $1.08 \times 10^{-4}$ |
| SMAD2⟷ESR1 | SMAD | Nuclear receptor | $3.61 \times 10^{-2}$ |
| SMAD3, SMAD4⟷ETS1, SMARCC2 | SMAD | Ets, Myb/SANT | $8.59 \times 10^{-3}$ |
| SMAD3, SMAD4⟷CTCF | SMAD | C2H2 ZF | $1.05 \times 10^{-3}$ |
| RFX4⟷RFX2 | RFX | RFX | $8.59 \times 10^{-3}$ |
| THRA, RXRB⟷NR2C2 | Nuclear receptor | Nuclear receptor | $8.59 \times 10^{-3}$ |
| ESR1⟷SP3 | Nuclear receptor | C2H2 ZF | $8.59 \times 10^{-3}$ |
| ESR1⟷KLF5, KLF2, KLF4 | Nuclear receptor | C2H2 ZF | $4.27 \times 10^{-3}$ |
| ESR1⟷ZBTB1 | Nuclear receptor | C2H2 ZF | $1.05 \times 10^{-3}$ |
| TFAP4⟷CTCF | bHLH | C2H2 ZF | $1.77 \times 10^{-2}$ |
| PURA⟷ESR2, ESRRA | Unknown | Nuclear receptor | $1.77 \times 10^{-2}$ |
| KLF8⟷CTCF | C2H2 ZF | C2H2 ZF | $4.93 \times 10^{-4}$ |
| HEY2⟷HAND1 | bHLH | bHLH | $3.61 \times 10^{-2}$ |
| KLF15⟷CTCF | C2H2 ZF | C2H2 ZF | $4.93 \times 10^{-4}$ |
| SP8, KLF14, KLF16⟷CTCF | C2H2 ZF | C2H2 ZF | $6.06 \times 10^{-7}$ |

**Table S16.** Significant matches with HIPPIE database inferred by SATORI deep with Pre-loaded CNN and Entropy Loss using Attn-Attr scores on Human Promoters Dataset

| TF Interaction | TF1 Family | TF2 Family | P-value |
|---|---|---|---|
| MBD2⟷ESR1 | MBD | Nuclear receptor | $5.53 \times 10^{-3}$ |
| NR0B1⟷ESR1 | Unknown | Nuclear receptor | $5.53 \times 10^{-3}$ |
| NR0B1⟷PPARG | Unknown | Nuclear receptor | $2.18 \times 10^{-2}$ |
| SP2, SP1⟷ESR1 | C2H2 ZF | Nuclear receptor | $5.53 \times 10^{-3}$ |
| SP2, SP1⟷ETS1, SMARCC2 | C2H2 ZF | Ets, Myb/SANT | $1.39 \times 10^{-3}$ |
| NFIA, NFIB, NFIX⟷SMAD2 | SMAD | SMAD | $2.18 \times 10^{-2}$ |
| NFIA, NFIB, NFIX⟷MYF5, MYOD1 | SMAD | bHLH | $4.31 \times 10^{-2}$ |
| JUN⟷PURA | bZIP | Unknown | $4.31 \times 10^{-2}$ |
| WT1⟷NR2C2 | C2H2 ZF | Nuclear receptor | $3.63 \times 10^{-5}$ |
| SMAD2⟷ESR1 | SMAD | Nuclear receptor | $4.31 \times 10^{-2}$ |
| TFCP2⟷PBX3, NFYC | Grainyhead | Homeodomain, Unknown | $1.09 \times 10^{-2}$ |
| SMAD3, SMAD4⟷ETS1, SMARCC2 | SMAD | Ets, Myb/SANT | $1.09 \times 10^{-2}$ |
| SMAD3, SMAD4⟷CTCF | SMAD | C2H2 ZF | $2.80 \times 10^{-3}$ |
| RFX4⟷RFX2 | RFX | RFX | $2.18 \times 10^{-2}$ |
| THRA, RXRB⟷NR2C2 | Nuclear receptor | Nuclear receptor | $1.09 \times 10^{-2}$ |
| ESR1⟷SP3 | Nuclear receptor | C2H2 ZF | $1.09 \times 10^{-2}$ |
| ESR1⟷KLF5, KLF2, KLF4 | Nuclear receptor | C2H2 ZF | $5.53 \times 10^{-3}$ |
| ESR1⟷ZBTB1 | Nuclear receptor | C2H2 ZF | $1.39 \times 10^{-3}$ |
| TFAP4⟷CTCF | bHLH | C2H2 ZF | $2.18 \times 10^{-2}$ |
| ETS1, SMARCC2⟷KDM2B | Ets, Myb/SANT | CxxC | $1.55 \times 10^{-4}$ |
| ETS1, SMARCC2⟷ETS2 | Ets, Myb/SANT | Ets | $6.59 \times 10^{-4}$ |
| PURA⟷ESR2, ESRRA | Unknown | Nuclear receptor | $2.18 \times 10^{-2}$ |
| KLF8⟷CTCF | C2H2 ZF | C2H2 ZF | $6.59 \times 10^{-4}$ |
| KLF15⟷CTCF | C2H2 ZF | C2H2 ZF | $6.59 \times 10^{-4}$ |
| SP8, KLF14, KLF16⟷CTCF | C2H2 ZF | C2H2 ZF | $9.43 \times 10^{-7}$ |

### REFERENCES

1. R. Z. Li, C. Z. Han, and C. K. Glass, "TIANA: transcription factors cooperativity inference analysis with neural attention," BMC bioinformatics **25**, 274 (2024).
2. A. Vaswani, N. Shazeer, N. Parmar, *et al.*, "Attention is all you need," Adv. neural information processing systems **30** (2017).
